# Episodic boundaries affect neural features of representational drift in humans

**DOI:** 10.1101/2023.08.20.553078

**Authors:** Nimay Kulkarni, Bradley C. Lega

## Abstract

A core feature of episodic memory is representational drift, the gradual change in aggregate oscillatory features that supports temporal association of memory items. However, models of drift overlook the role of episodic boundaries, which indicate a shift from prior to current context states. Our study focuses on the impact of task boundaries on representational drift in the parietal and temporal lobes in 99 subjects during a free recall task. Using intracranial EEG recordings, we show boundary representations reset gamma band drift in the medial parietal lobe, selectively enhancing the recall of early list (primacy) items. Conversely, the lateral temporal cortex shows increased drift for recalled items but lacked sensitivity to task boundaries. Our results suggest regional sensitivity to varied contextual features: the lateral temporal cortex uses drift to differentiate items, while the medial parietal lobe uses drift-resets to associate items with the current context. We propose drift represents relational information tailored to a region’s sensitivity to unique contextual elements. Our findings offer a mechanism to integrate models of temporal association by drift with event segmentation by episodic boundaries.

## Introduction

A defining feature of episodic memory is the contextual information that associates and differentiates item representations. Temporal relationships are a key component of contextual information (***Tulving and Markowitsch, 1998***; ***Fortin et al., 2002***; ***Sederberg et al., 2008***). The successful encoding of these temporal relationships has been tied to a drifting pattern in neural activity (***Folkerts et al., 2018***; ***Mau et al., 2020***). However, temporal relationships are also influenced by large context shifts, called boundaries (***Zacks et al., 2001, 2007***; ***DuBrow and Davachi, 2013***; ***Ranganath, 2019***). On an intuitive level, relational information may be encoded through both mechanisms: association with preceding items, represented by drift, and the association with shifts in context, represented by boundaries. However, it remains unclear how a drifting signal, characterized by gradual changes neural activity, accommodates large shifts in context represented by boundary signals. The primary focus of this study is to reconcile these mechanisms of context and clarify how the brain simultaneously represents item-specific temporal information and item-independent boundary information to form a rich contextual background to encode new memories.

The Temporal Context Model (TCM) suggests an item’s temporal context is built from items that immediately precede it (***Howard and Kahana, 2002***). When a new item is introduced, it updates the context state and gradually changes the contextual representation. The degree of item similarity *drifts* such that items presented close together have high contextual overlap while items far apart have less overlap. This drifting context offers a mechanism for robust behavioral effects observed in memory.

Drift in context representation provides a compelling explanation for the recency and contiguity effects. The recency effect arises from the proximity of recency events to the recall period. The temporal context state at the recall period has not drifted significantly from the context state while encoding the recency items (***Lohnas et al., 2015***); therefore, items presented most recently share the highest similarity to the context state at retrieval allowing recency items to be most easily remembered. While retrieving a specific event, the recollection process not only reinstates that item’s encoding representation but also reinstates preceding item representations as context (***Manning et al., 2011***). Reinstated context serves as a cue mechanism for nearby items resulting in tendency to cluster items presented together, referred to as the contiguity effect (***Folkerts et al., 2018***).

However, these models do not inherently capture primacy effects as convincingly, which have been hypothesized to arise from rehearsal (***Tan and Ward, 2000***) or less well understood processes such as neural fatigue (***Serruya et al., 2014***; ***Lohnas et al., 2020***). Updated versions of TCM model primacy using terms that lack an obvious neural correlate (***Polyn et al., 2009***; ***Lohnas et al., 2015***) creating a need for a coherent integration of primacy and drifting information for context representation.

Boundary-related information provides a potential mechanism for understanding primacy. In parallel with a gradually drifting context representation, episodic experience is also segmented by sharp shifts in context, known as boundaries. Boundaries are created after large changes in stimuli, task-goals, or context information (***Kurby and Zacks, 2008***; ***Baldassano et al., 2017***; ***Griffiths and Fuentemilla, 2019***). Event segmentation theory holds that boundaries segment continuous experience into “events”, resulting in improved memory within the current event but worsened memory of a prior event. Boundaries may exert this effect by disrupting contextual associations between items separated by a boundary (***DuBrow and Davachi, 2013***; ***Ezzyat and Davachi, 2014***). Conversely, random boundary insertions improve memory of subsequent items and induce a novel primacy-like effect (***Mau et al., 2020***; ***Ranganath, 2019***). The ability of boundaries to disrupt associations of prior items but improve associations of subsequent items suggests they may play a role in defining context. However, the effect of boundaries on a drifting context signal is still unclear. Ranganath and colleagues suggest boundaries reset a portion of temporal context and drift occurs relative to a reinstated boundary signal (***Pu et al., 2022***). Alternatively, boundaries may accelerate the rate of drift inducing a greater contextual dissimilarity between boundary separated items (***Horner et al., 2016***; ***DuBrow et al., 2017***).

Strong evidence supports the presence of boundary representations within the medial temporal regions. Aggregate regional activity (***Baldassano et al., 2017***) and individual neurons (***Yoo et al., 2022***; ***Zheng et al., 2022***) show sensitivity to embedded boundaries in several tasks of episodic memory. However, boundary representations are not confined to just the MTL, studies across multiple cognitive domains highlight their presence in the parietal lobe. In studies of spatial navigation, boundary vector neurons are specifically sensitive to spatial boundaries (***Alexander et al., 2020***). These spatial boundaries help create a generalized map of the surrounding area (***Bicanski and Burgess, 2020***). Similarly, in episodic tasks, parietal regions demonstrate content-invariant activation at conceptual boundaries (***Zacks et al., 2001***; ***Baldassano et al., 2017***). These boundary representations play a key role in the construction of situational models (***Kurby and Zacks, 2008***). Whether spatial or episodic, the parietal lobe’s ability to represent boundaries may underlie its role in constructing a abstract contextual framework as described by the posterior medial-anterior temporal (PMAT) framework (***Ranganath and Ritchey, 2012***). With these (non-invasive) findings as motivation, we sought to analyze neural evidence of drift-like and boundary related information recorded from temporal (hippocampus and lateral temporal cortex) and medial parietal (precuneus and posterior circulate cortex) regions in 99 human intracranial EEG patients. We find recalled items drift at a higher rate than non-recalled items. Across multiple lists, boundary items share significant spectral similarity mainly in the parietal lobe. Our findings suggest that these regions provide complementary representations of temporal context and boundary-related activity, with medial parietal areas exhibiting unique sensitivity to boundary-related information linked with memory performance. Our data align with experiments in rodents that link parietal activity to boundary-representations in spatial navigation, suggesting a more generalized role of the parietal regions in event segmentation which supports multi-scale contextual associations in episodic memory.

## Results

We analyzed depth electrode recordings from participants performing the delayed free recall task (Figure 1A). The task’s robust contiguity effect as well as its predictable internal structure allowed us to study both item-boundary and item-item relationships. Participants had an overall recall rate of 24% ± 11% (mean ± std; Figure 1D). Our analysis focused on gamma band (30 to 100 Hz) power from electrodes in the medial parietal lobe (MPL; precuneus and posterior cingulate cortex), the lateral temporal cortex (LTC; superior and medial temporal gyri), and the hippocampus (HC). We focused on gamma power due its strong association with single neuron activity (***Umbach et al., 2020***), local excitatory-inhibitory feedback loops (***Buzsaki and Wang, 2012***; ***Wang, 2010***), and orde. These regions were chosen because structural and functional studies suggest the posterior medial (PM) and anterior temporal (AT) regions represent different features of episodic memory that are integrated by the hippocampus (***Ranganath and Ritchey, 2012***). Additionally, these regions have robust electrode coverage in patients intracranial EEG monitoring. To evaluate the relatedness of item representations, we used principal component analysis (PCA) to identify common features of gamma band covariance and quantified the similarity of between items with cosine similarity as a function of positional distance between item pairs, see Methods.

**Figure 1.**
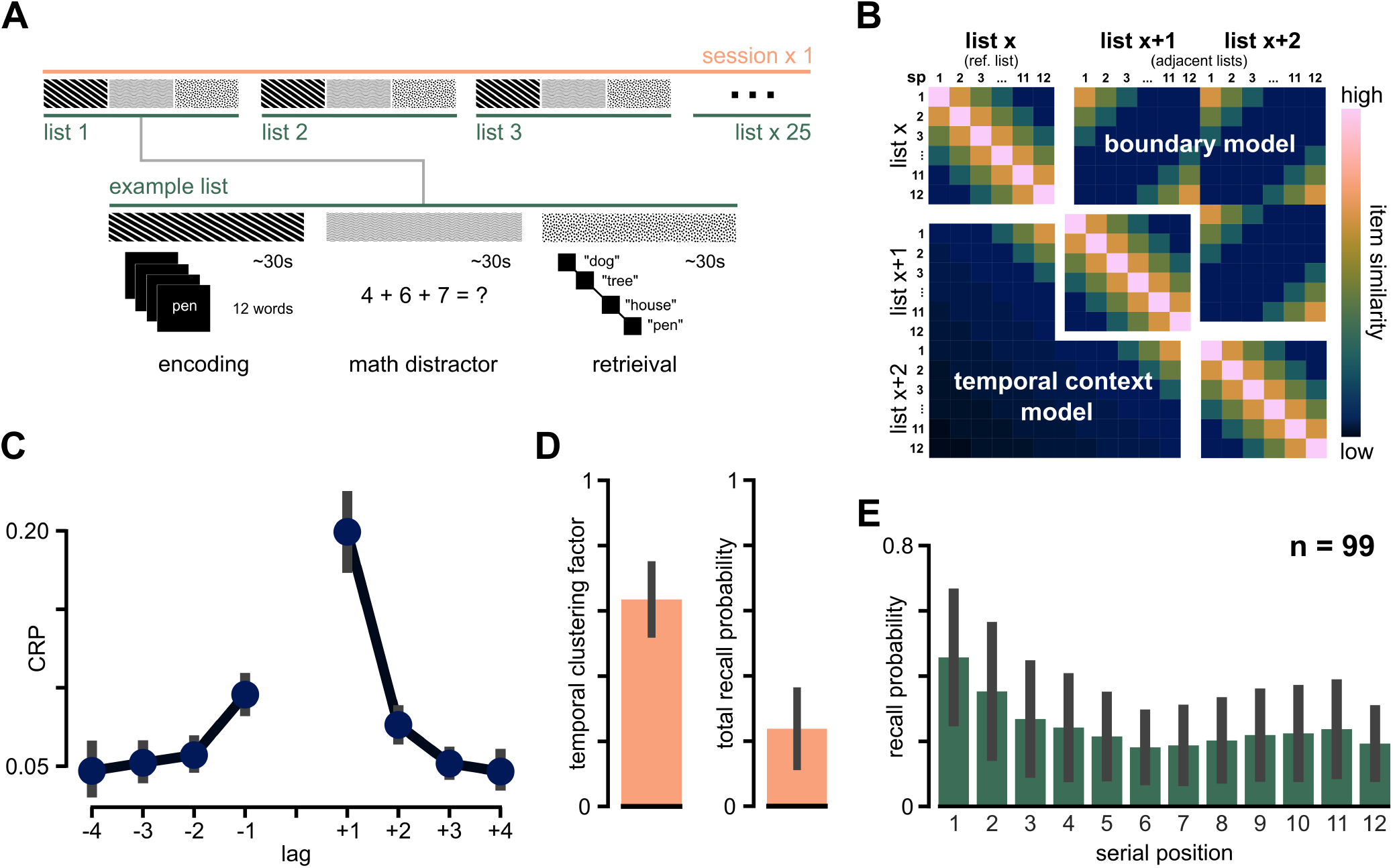
Illustration of task and behavioral summary. **A**. Diagram of the delayed free recall task. **B**. Schematic of the predicted similarity between items across in the same list (*list x*) and 2 adjacent lists (*list x+1, x+2*). Items presented close together in time show the most representational similarity, shown by the drifting reduction in similarity perpendicular to the diagonal. TCM predicts item representations should continue to decay across lists due to increased temporal distance shown as blue transitions to black (bottom left). A boundary-oriented representation would show similarity between boundary items across multiple lists (top right). **C**. The average proportion of recalled events for each serial position for each participant (n=99). **D**. The average recall probability (left, n=99, mean±sd: 0.24 ± 0.11) and the average temporal clustering factor (TCF; right, 0.64 ± 0.10) of all study participants. **E**. The lagged conditional response probability (CRP) is shown for 4 lags in either direction characterizing the contiguity in our data. The error bars in (E) represent ±1SEM. Error bars denote ±1SD unless otherwise noted.

### Enhanced drift rate increases distinctiveness of spectral representations for recalled items

First, we focus on neural similarity between words in the same list. Previous work consistent with temporal context models suggests neural pattern similarity reduces as a function of distance between related memories (***Hsieh et al., 2014***; ***Hsieh and Ranganath, 2015***; ***El-Kalliny et al., 2019***). In concert with previous work, we refer to this phenomenon as neural drift. To quantify drift in our data, we calculated a similarity value between every item in the same list (Figure 2A).

**Figure 2.**
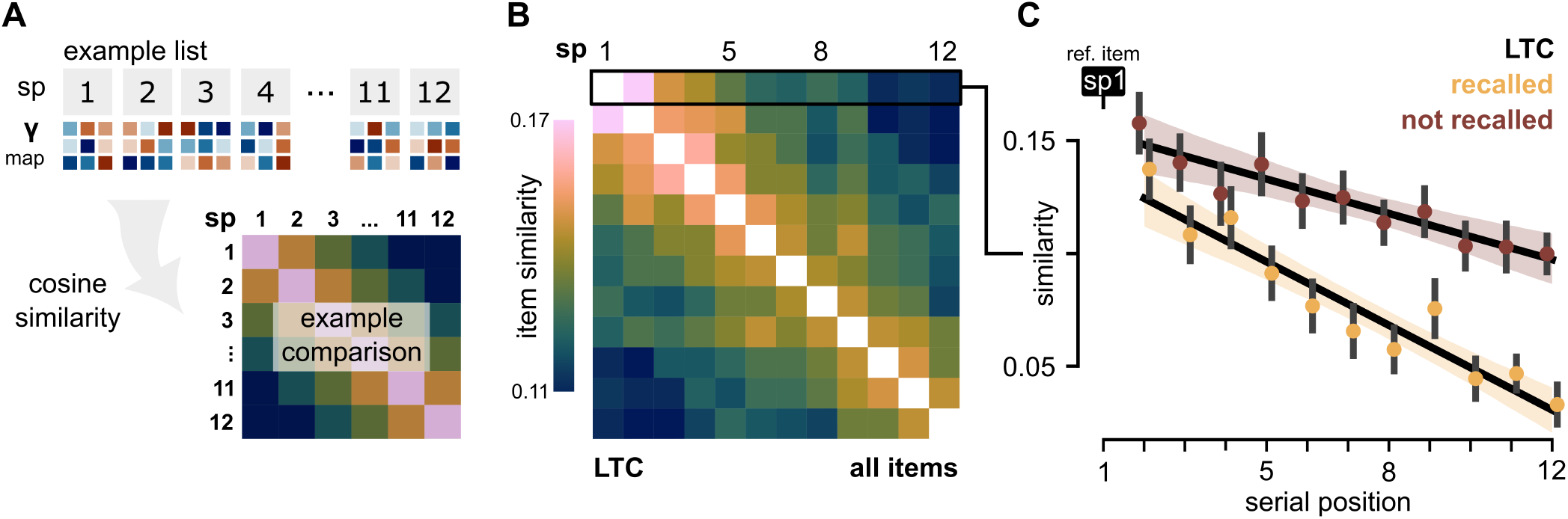
Neural drift is enhanced between recalled item pairs within the lateral temporal cortex. **A**. Schematic of similarity analysis between item pairs in the same list. (sp: serial position; *γ* map: gamma spectrum, PCA-optimized, time-frequency map) **B**. Similarity of all item pairs regardless of recall status, averaged by subject, across all electrodes in the LTC. **C**. Similarity of serial position 1 to each subsequent item in a list contrasting when both items are recalled (yellow) to when both items are not recalled (red). The error bars demonstrate variation in average similarity between participants. The trend lines are for visualization purposes only, the LME used to statistically test the result is described in the results section. Error bars indicate ±1 SEM. **Figure 2—figure supplement 1**. Drift rate does not differ between recalled and non-recalled items in the hippocampus and lateral temporal cortex

Figure 2B shows the average similarity of every pair of items in a list for electrodes in the LTC. We found item similarity was reduced with increasing inter-item distance, aligning with prior findings (***Hsieh et al., 2014***; ***El-Kalliny et al., 2019***). However, words at different serial positions are recalled at different rates. To understand how item-item similarity is affected based on successful recall, we compare item pairs where both items are recalled to pairs where both are not recalled (2C) using a linear mixed effects model (LMEM). The LMEM compared similarity of each serial position (SP) in reference to the first item with an interaction term for recall success, see Methods. In the lateral temporal cortex, we found that similarity decreased with increasing inter-item distance and that recalled pairs tended to be less similar than non-recalled pairs (LMEM, n=81; *β*_SP_=-0.005, CI:[-0.007, -0.003], *p*_SP_ < 0.001; Interaction: SP x recalled items=-0.004, [-0.007, -0.002], *p*_interaction_ = 0.001). In terms of drift, items drifted at a higher rate such that individual item representations were less similar, or more distinct, when participants recalled both items.

### Strong boundary representation in medial parietal lobe augments recall of initial list items

Next, we were interested in how boundaries affect item representations. In our behavioral task, the beginning of each list serves as a consistent boundary marking the initiation of a new encoding period. We hypothesized that a boundary signal would be most strongly represented in the first item of each list and its representation would be similar across multiple lists.

To test this hypothesis, we extended the item-item comparisons described in Figure 2 to compare items *across adjacent lists*. The schematic in Figure 1B visualizes the predicted similarity between items in the reference list (denoted list x) and multiple adjacent lists (denoted list x+1 and x+2). We contrast two possible models of association in the same schematic. Models based on continuous effects of new items on neural similarity (namely, the temporal context model) suggest that item-item associations would show decreasing similarity with increased positional distance, even across lists, without any evidence of enhanced similarity across lists for boundary-adjacent items. An alternative model that makes distinct allowance for boundary information would show evidence of recurring activity features near boundaries, (ie. the beginning and end of the item lists) with consistently decreasing similarity across lists for middle-list items. The schematic provides a visualization of our predictions captured in the models we applied to neural data, described below.

We began by calculating similarity between serial position (SP) 1 and every item in the same list (*list x*) and the 3 adjacent lists (*list x+1, x+2, x+3*), shown in Figure 3A. The similarity of SP1 items compared to each item in the adjacent lists for the medial parietal lobe, averaged by subject, is shown in Figure 3E. To calculate the degree of boundary representation, we built a general linear model with two predictor variables: boundary proximity, the distance of an item to the beginning of the list, and list distance, the distance of the target list from the reference list. An example subject’s averaged similarity data is shown in Figure 3D with the model-fit superimposed. We fit the GLM to the average similarity of *list x, SP1* to every item in list x+1, x+2 and x+3 for each subject individually in each region of interest. We found SP1s across adjacent list were most similar in the medial parietal lobe as compared to the lateral or medial temporal cortex (Figure3B; t-test: MPL-LTC, t(83,81)=4.05, p=8.0 × 10^−5^; MPL-HC, t(83,76)=3.27, p=1.3 × 10^−3^). The similarity of *list x, SP1* to subsequent boundary items in these regions is less pronounced (3–figure supplement 1 A,C). We repeated the same analysis to contrast recalled item pairs with non-recalled item pairs to assess the effect of recall success on boundary representation. We observed the similarity of boundary items *across item lists* was higher for recalled pairs (Figure 3B, right; paired t-test, t(83)=2.45, p=0.02). This finding suggests an elevated boundary representation is incorporated for strongly encoded items, and it stands in contrast to lower similarity observed for recalled items across item lists in the LTC and HC. Finally, we asked whether the strength of boundary similarity correlated with the proportion of successful SP1 recalls per participant. We found participants with stronger boundary item similarity in the medial parietal lobe tended to recall a greater proportion of presented SP1 items (linear regression, r=0.41, t(83)=4.08, p=1.1 × 10^−4^).

**Figure 3.**
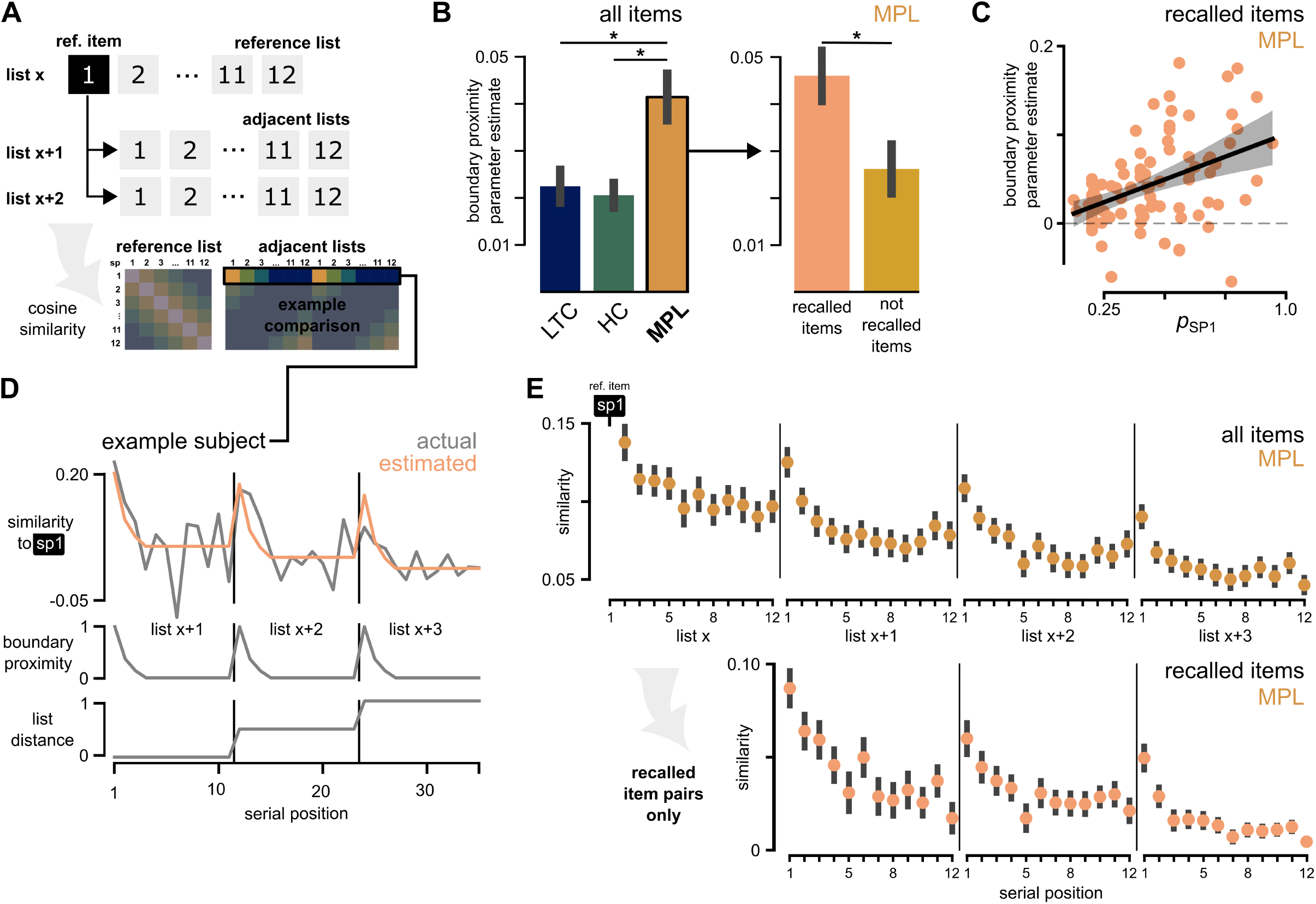
The first items (SP1) across adjacent lists share a boundary representation. **A**. A schematic showing the comparison of SP1 to every item in the adjacent three lists. **B**. The mean parameter estimate of the predictor for boundary proximity contrasted by region for all item pairs (left; LTC: n=81, mean β=0.020; HC: n=76, mean β=0.022; MPL: n=83, mean β=0.042). The parameter estimate of recalled-only pairs and non-recalled pairs is also shown for the MPL (right; recalled: pink, mean β=0.045; non-recalled: orange, mean β=0.027). **C**. Correlation of proportion of recalled SP1 by subject to the parameter estimate for boundary proximity (ordinary least squares regression, r=0.41, p=1.1 × 10^−4^. **D**. Design of the GLM used to fit adjacent list similarity in the MPL. The response variable (similarity to *list x, SP1*; gray, top) shown for an example subject with the model prediction (orange, top) overlayed. The predictor variables, boundary proximity (middle) and list distance (gray, bottom), are also shown. **E**. Mean similarity of all serial positions to the reference item (*list x, SP1*) regardless of recollection (top). Below, the mean similarity of only instances where both *list x, SP1* and the target item are recalled are shown for all serial positions in adjacent lists (*list x+1, x+2, x+3*; bottom). The error bars demonstrate variation in average similarity between participants. All error bars indicate ±1 SEM. ∗ *p* < 0.05 **Figure 3—figure supplement 1**. Serial position 1 does not show boundary representation in the hippocampus and lateral temporal cortex. **Figure 3—figure supplement 2**. Boundary information is represented in multiple principal components

### The boundary representation is conserved in multiple primacy events

Our analysis thus far of adjacent SP1 items supports the presence of conserved boundary representation apart from drift-like changes in neural activity. This pattern is specific to the MPL suggesting this region represents item context relative to a boundary. To further investigate the role of boundary information in episodic encoding, we tested whether neural activity for boundary-adjacent serial positions (ie. SP2 and SP3, items which are subsequent to the boundary item within an encoding list) exhibited persistent boundary information using the same cross-list analysis as described above.

We modified our prior analysis to test the similarity of *list x, SP2 and SP3* to every item in the adjacent three lists for recalled item-pairs only (Figure 4A). We tested for boundary representation in these serial positions to distinguish the role of the medial parietal lobe in representing contextual features. If the MPL is specifically sensitive to boundary information, SP2 and SP3 should show greater similarity to the boundary item than any other items in the same list. In support of our hypothesis, we found both SP2 and SP3 shared more similarity to the adjacent lists’ SP1 rather than the adjacent lists’ SP2 or SP3, respectively (Figure 4). We tested this by comparing the average similarity per subject of *list x, SP2* to SP2 in each of the 3 adjacent lists. For SP2, we found SP2 was more similar to SP1 than SP2 in list x+3 (4C, top; two-sided, paired t-test, t(83)=2.98, p=0.003). We also found SP3 from list x was more similar to SP1 in both list x+2 and list x+3 than to SP3 in the same list (4C, bottom; list x+2: t(83)=2.49, p=0.015, list x+3: t(83)=2.31, p=0.023). In summary, multiple items in primacy positions share a strong similarity to boundaries in other lists. Importantly, this cross-list similarity appears to be greater than the degree of similarity to their own serial position in other lists. We examine the potential implications for this finding more thoroughly and within the temporal context framework (versus other serial encoding models) in the Discussion section.

**Figure 4.**
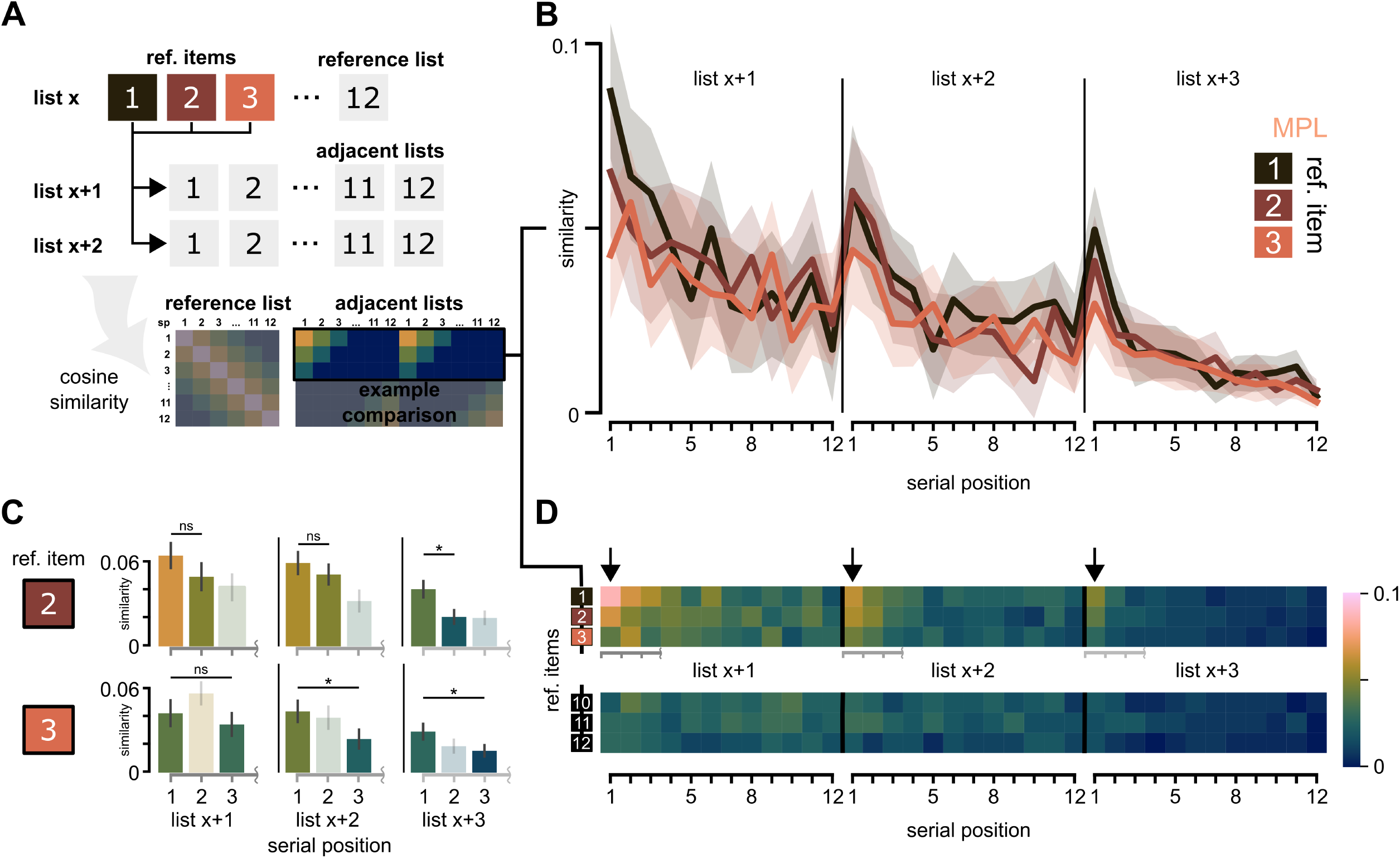
Multiple primacy items retain similarity to the boundary representation in the MPL. **A**. A schematic of showing comparison of the reference and target items (top). The highlighted portion of the matrix demonstrates hypothesized trend in the boundary model presented as a part of 1B (bottom). **B**. Averaged similarity of only recalled item-pairs comparing the reference items: *list x, SP1-3* to each item in *list x+1, x+2, x+3*. A representation of boundary would be supported by *list x, SP2* showing more similarity SP1 in the adjacent lists rather than its own serial position. Alternatively, positional coding, the similarity of the same serial position between adjacent lists, would show greater similarity between *list x, SP2* and SP2 in adjacent lists. Note the similarity of *list x, SP2 and SP3* to SP1 in list x+3 rather than SP2 and SP3 in list x+3. The confidence interval demonstrates ±1SEM of variation in average similarity between participants. **C**. Comparison of the similarity between reference items: *list x, SP2 and SP3* to the first three serial positions in the adjacent lists. The bar graphs demonstrate the similarity between SP2 and SP3 to the boundary item compared to the same serial position in the corresponding list. The bar colors reflect the magnitude of similarity as shown in (D). **D**. Same as (B) shown in matrix form for *list x, SP1-3 and SP10-12* to demonstrate the lack of boundary similarity of end-list items. The black arrows indicate the first serial position (SP1) of the adjacent lists and emphasize the presence of boundary representation. The similarity of *list x, SP2 and SP3* to the boundary is statistically tested in (C) (gray axes). All error bars indicate ±1SEM.

### Non-recalled items show increased boundary representation at the end of the list

Our results so far support the presence of a recurring boundary signal at the beginning of each list. Next, we were interested in whether a similar boundary representation existed at the end of each list. We repeated the same analysis described in Figure 3 but for *list x, SP12* to each item in the adjacent lists (Figure 5A). When comparing the similarity of all items pairs, we found later serial positions in adjacent lists shared a gradually increasing similarity to SP12 (Figure 5B). However, when we filtered for recalled items only, this trend disappeared (5D, pink). When recalled items from *list x, SP12* were compared to adjacent recalled items, the items did not demonstrate any position dependent pattern. To test this trend, we fit a similar GLM to Figure 3D, with boundary proximity as a linear predictor (see Figure 5–figure supplement 1C), which showed a significantly higher boundary parameter estimate for non-recalled item pairs (Figure 5C, right; paired t-test, t(74)=-3.39, p=1.1 × 10^−3^). These findings suggest end-list items are represented more distinctly, or less similarly, to all succeeding items. Stated another way, while our analysis identifies persistent boundary-information across items lists for both beginning of list (primacy) and end of list (recency) items, these two classes diverge in terms of the apparent connection to recall success. Non-recalled items tended to show higher boundary similarity at the end of the list. Conversely, recalled items showed greater boundary similarity at the beginning of the list but not at the end of the list. Taken together, the greater preservation of boundary-related information seems to support recall for primacy items but not recency items.

**Figure 5.**
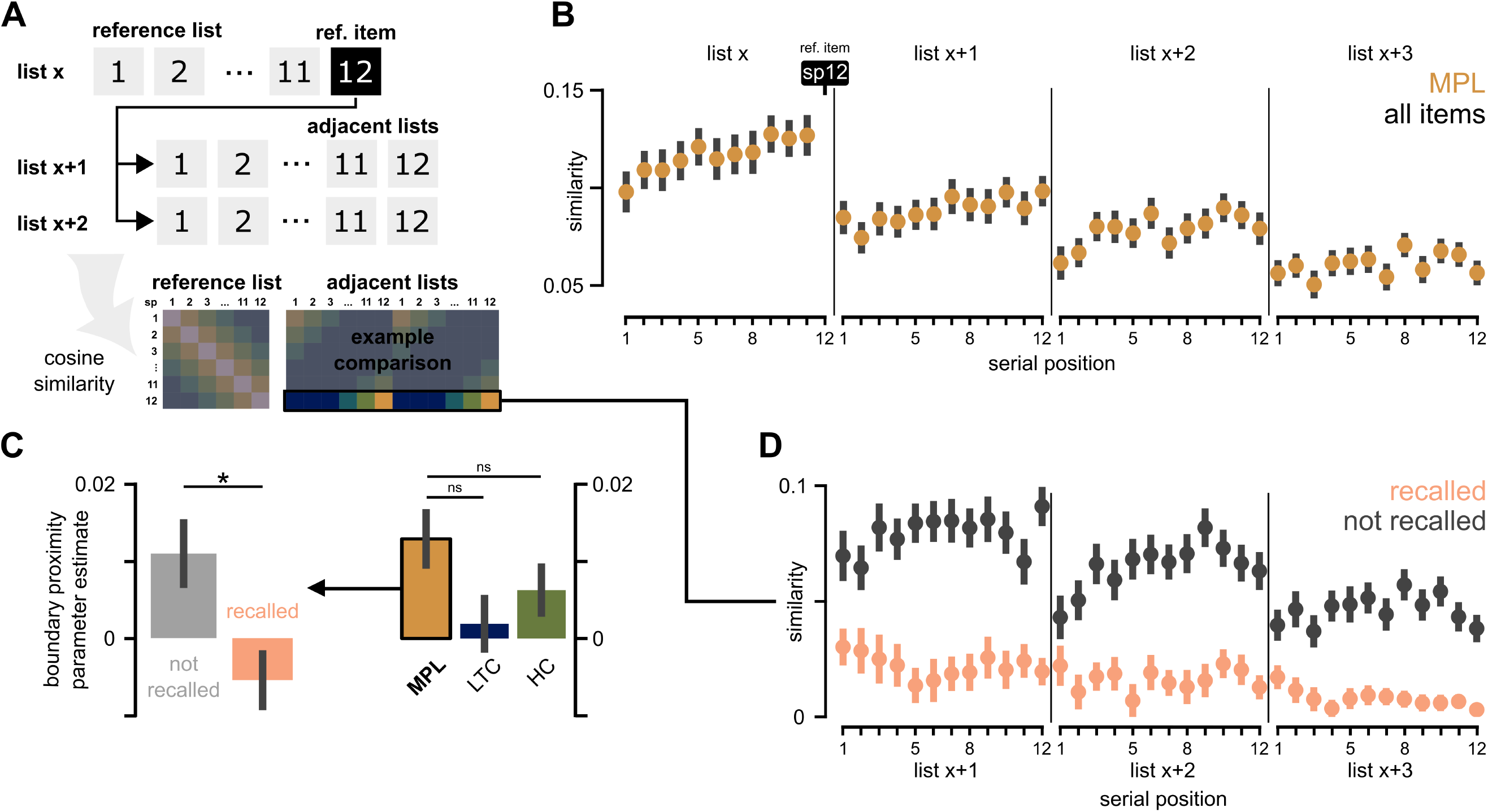
The last item shares similarity to other items at the end of adjacent lists in the MPL. **A**. Schematic of the predicted similarity between the last item in a list *list x, SP12* to each item in the adjacent lists. **B**. Mean similarity of all serial positions to the reference item (*list x, SP12*) regardless of recollection (top). This shows the similarity of end-list items to other end-list items across adjacent lists. Importantly, the last item in each list (SP12) is not similar to SP1 of the adjacent list despite the temporal proximity. The error bars demonstrate ±1SEM of variation in average similarity between participants. **C**. The mean parameter estimate of the predictor for boundary proximity fit to average similarity data from the MPL for recalled versus non-recalled items (left; recalled: pink, mean β=-0.006; non-recalled: orange, mean β=0.012). Additionally, the mean parameter estimate for all items between regions. (Right; MPL: n=83, mean β=0.011; LTC: n=81, mean β=0.005; HC: n=76, mean β=0.003). **D**. Same as **B** for recalled pairs only. All error bars indicate the ±1SEM. **Figure 5—figure supplement 1**. End-list items show partial boundary similarity to end-list items across multiple regions

## Discussion

In the present study, we explored the role of task boundaries in shaping the contextual representation in episodic memory. In a delayed free recall task, we used the beginning and end of the list as repeated boundaries to investigate their influence on drift, specifically examining regional variation. In the medial parietal lobe, we discovered items adjacent to boundaries shared remarkable similarity across multiple lists, indicative of a recurring boundary signal. The strength of the boundary signal correlated with the successful recollection of boundary items. Next, we show the boundary signal is also present in serial positions two and three items removed from the boundary, further supporting the role of a boundary as a key feature of (associative) context for these items. Finally, we demonstrate evidence of recurring feature similarity for end of list boundaries (recency). However, unlike primacy items, we did not observe enhanced similarity for successfully recalled items raising the possibility that recency boundaries interfere with (rather than enhance) encoding processes. Overall, our study suggests a regional sensitivity to distinct contextual elements which, depending on the serial position, can variably affect item recall.

We began by presenting evidence that drift in the gamma spectrum is a memory relevant phenomenon. Our findings corroborate existing evidence suggesting neural drift is a marker of association between related memories (***Howard and Natu, 2005***; ***Buzsaki and Tingley, 2018***). Previous investigations found drift in the lateral temporal context contributed to successful recall of related items (***El-Kalliny et al., 2019***). Gamma oscillations also coordinate contextually relevant neuronal assemblies (***Umbach et al., 2022***). Our results build on this finding by characterizing drift in gamma oscillations and further describing how gamma activity is associated with context.

We also found a higher rate of gamma signal drift between successfully recalled items. This elevated drift rate might allow the representations of recalled items to remain distinct but ordered in memory. These findings align with several previous studies describing drift in item representations only when temporal order information is preserved (***Hsieh et al., 2014***; ***Hsieh and Ranganath, 2015***; ***Howard and Eichenbaum, 2015***). More generally, our findings support the mnemonic relevance of high frequency oscillations.

Next, we asked how gamma spectral activity reflects contextual association between items. In the medial parietal lobe, we observed recurring similarity between items distant in time but adjacent to boundaries. This pattern suggests spectral activity may carry information about an item’s relationship to a boundary. These observations align with the Context Maintenance and Retrieval model (***Polyn et al., 2009, 2012***) which extends the predictions of TCM to encompass broader relationships among items. Our results specify the spectral and regional properties of boundaryrelated features and demonstrate boundaries as an important aspect of context in episodic memory.

Our main finding shows specific similarity of serial position 1 across multiple lists (Figure 3). No-tably, this position lacks a preceding item, leading us to propose that the context for the first item may be the boundary itself. The behavioral association between boundary-related features in medial parietal cortex and recall fraction for serial position 1 items further supports the importance of this information in memory. Then, in accordance with TCM, the context for the second item would be derived from the first item, supported by the pronounced boundary signal in SP2 and SP3 (Figure 4). These results suggest that boundary representation might hold a critical role in primacy items specifically. Previous work also supports the idea of boundaries as drivers of primacy memory (***Pu et al., 2022***; ***Polyn et al., 2012***). Pu and colleagues introduced random boundaries in a task of ordered memory and reported an enhanced recollection of items immediately after the boundary, which they coined a local-primacy effect. Taken together, boundary representation may hold a critical role in consolidating primacy items in memory, introducing a new facet to our understanding of the primacy effect.

Conversely, an opposite effect of boundary representation was present for end of list items, showing an unexpected relationship of the end-list boundary with recall success. We found spectral similarity among these items when they are not recalled. Yet, when an end-of-list item is recalled, the item shares minimal similarity to succeeding items. This suggests distinctiveness is more important for recency items than their association with a boundary. Stated another way, for recency items, drift-like features consistent with predictions of the temporal context model may principally provide a cue for recall rather boundary information. Our findings suggest a broader scope of contextual association than just prior items, where temporal proximity as well as task structure in the form of boundaries, play intertwined roles in contextual construction. Our data therefore have implications for updated iterations of the temporal context model incorporating (perhaps) specific terms for boundary information. This may in turn provide a more systematic prediction of primacy effects in behavioral data.

Next, we propose drift is governed by varying contextual features. In the medial parietal lobe, spectral activity aligned closely with task boundaries. Drift between item pairs seemed to reset at each boundary, leading to renewed similarity after each boundary. This observation aligns with previous work suggesting boundaries reset temporal context (***Pu et al., 2022***). In the temporal cortex, our findings extend prior studies which suggest the temporal lobe may play a role in associating adjacently presented items (***Yaffe et al., 2014***; ***El-Kalliny et al., 2019***). We found items encoded in distant serial positions, but within the same list, drifted significantly more than items from nearby serial positions (Figure 2C). Consistent with the predictions of the temporal context model, the reduced similarity between distant items may reflect reduced contextual overlap pro-portional to the time elapsed between them. However, across task boundaries, our study did not detect a robust change in drift pattern in the medial or lateral temporal cortex. This finding contrasts with significant work (***Ben-Yakov and Henson, 2018***; ***Ezzyat and Davachi, 2014***; ***Griffiths and Fuentemilla, 2019***) which shows hippocampal sensitivity to event-boundaries. One interpretation would be that boundary representations in the hippocampus are quite sparse and represented by populations of time-sensitive cells whose activity is indexed to task-related boundaries (***Umbach et al., 2020***). While the sparse representations may not be detectable in gamma activity, perhaps it suggests drift in these regions represents a more abstract set of contextual features accumulated from multiple brain regions (***Baldassano et al., 2017***).

Finally, we build on the idea of boundary-modulated drift by proposing region-specific drift rates that occurs relative to distinct contextual features. The presence of boundary representations in the medial parietal lobe aligns with prior suggestions that the posterior medial network plays a role in constructing situational models (***Ranganath and Ritchey, 2012***). This differential sensitivity to contextual features may underlie more complex, multi-dimensional contextual associations. In free recall, temporal relationships are thought to dominate. However, our work suggests the incorporation of an additional, boundary-oriented feature of context plays an equally crucial role in organizing episodic memory.

## Methods and Materials

### Participants

This study included 99 human adult subjects (age: 39.0±11.1 years old, mean±sd; 45 female) from the University of Texas Southwestern Medical Center (UTSW) who underwent intracranial electroencephalographic (iEEG) electrode implantation for drug-resistant epilepsy. Electrode locations were chosen by a clinical team for seizure mapping. The UTSW institutional review board approved this study and each participant provided informed consent.

### Experimental design

Participants performed a delayed free recall task. Each session consisted of 25 lists of 12 common nouns, selected at random without replacement (http://memory.psych.upenn.edu/WordPools). During the encoding phase, each item in a list was shown for 1.6 seconds interspersed with a 1-second cross between events followed by a 30 second arithmetic distractor task. Participants were then asked to recall as many words as possible from the immediately preceding list in a 30 second retrieval period.

### Intracranial recordings

iEEG signal was recorded continuously for all implanted electrodes (Ad-tech or PMT depth electrodes, 0.8 mm) at 1000Hz for an entire session on a Nihon–Kohden 2100 clinical system. Contacts were localized by manual review by a clinical radiologist. The set of included participants included in this study had at least 1 electrode in the lateral temporal cortex (LTC; superior and medial temporal gyri), medial parietal lobe (MPL; posterior cingulate cortex and precuneus), or hippocampus (HC). Any electrodes in epileptogenic tissue or showing inter-ictal patterns were excluded from the dataset. Continuous iEEG data was time stamped at the beginning of each encoding event, distractor task, and retrieval event. Only encoding data, the 1600ms during which the word was on-screen, was considered in subjects who recalled at least 20 words in a session.

### Power and similarity analyses

Signal pre-processing and oscillatory power analyses were performed with MATLAB (The Math-Works, Inc., Nattick Massachusetts, United States) and Python using a combination of custom scripts and established EEG analysis toolboxes. EEG signals were downsampled to 500Hz and filtered using a band-pass filter between 2Hz and 100Hz. The spectral power was calculated by convolving with a complex Morlet wavlets (wave number = 6) for 10 linearly spaced frequencies in the gamma band (30-100Hz) excluding 1 frequency (63Hz) due to its proximity to 60Hz line noise were included in further analysis. Spectral data was log-transformed and z-scored per frequency in reference to baseline activity (-450ms to -250ms prior to item presentation). These methods align closely to established pre-processing pipelines (***Long et al., 2017***; ***Tan et al., 2020***; ***Cohen, 2014***).

The gamma time-frequency matrix was collapsed into a feature vector per event for principal component analysis (PCA) in MATLAB. PCA was applied to all encoding events per electrode. The 3 principal components with the highest explained variance were then used to reconstruct the power spectrum in time-frequency space (***Buzzell et al., 2022***). Each item-item comparison described below was calculated using cosine similarity according to the formula:

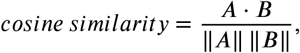

where *A* and *B* are the PCA-reconstructed gamma signal collapsed into a 1-dimensional time-frequency vector for the entire encoding duration of an event (1.6 seconds).

### Within list and across lists analyses

We conducted an item-item similarity analysis with a focus on change in similarity with increasing distance between items. We refer to a reference item within a reference list (see 3A) and compare that item’s spectral map to subsequent items, which we refer to as the target item in a target list. In Figure 2, each item in a reference list, denoted *list x*, was compared to every other item in the same list. In Figures 3 to 5, each reference item was also compared to the items in presented in the next 3 lists, denoted *list x+1, x+2, or x+3*. Similarity values of item-item comparisons were averaged within electrodes in the region of interest, then within each subject per serial position.

### Statistical analysis

In our within-list analysis (Figure 2), we employ a mixed effects model to examine the relationship between serial position and similarity amongst items in the same list. The fixed effects in the model were serial position, ranging from 1 to 12, and recall status, included as a binary categorical variable, to capture the interaction between serial position and successful recollection. Random nested terms for each electrode and subject were included to account for group level heterogeneity.

Across Figures 3 to 5, we used a General Linear Model (GLM) to approximate the relationship between the response variable: cosine similarity between item pairs of increasing inter-item distance, and two predictor variables: boundary proximity and list distance. The goal was to asses the impact of positional distance between items on the spectral similarity of the items. Boundary proximity captured the proximity of each serial position to the beginning or end of a list. For SP1, boundary proximity was quantified according to the equation:

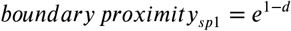

where *d* is the serial position of the target item within its respective list (Figure 3C). For SP12, we noted a linear increase in similarity for end-list events. Therefore, the boundary proximity predictor was defined as:

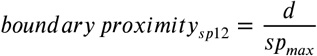

where *d* is the serial position of the target item in its respective list and *SP*_*max*_ is the number of items in each list (Figure 1C). The second predictor variable, list distance, quantified the overall decrease in similarity of distant lists. Items in list x+1, x+2, and x+3 were assigned values of 0, 0.5, and 1 respectively. The described GLM was fit to each subject’s averaged positional similarity individually and parameter estimates were compared using a unpaired, two-sided t-test between the parameter estimates for each subject across two regions of interest in Figures 3B and 5C.

### Behavioral metrics

We calculated temporal clustering factor (TCF) according to previously published methods (***Polyn et al., 2009***). For all retrieved words in a list, we determined the encoding serial position. We calculated the difference in encoding position for successively recalled items. If the first two recalled words were from encoding position 1 and 4, the difference would be 3. If the third recalled word was encoded in position 2, the next transition would be -2 (SP4 to SP2). Each transition received a score based on its rank amongst all remaining possible transitions. This metric yields a value per list which was averaged by session to generate an aggregate score of temporal clustering tendency for every subject.

The lagged conditional response probability (CRP) was similarly calculated according to prior studies (***Polyn et al., 2009***; ***Healey et al., 2019***). The CRP value at each lag is calculating dividing the counts of actual transitions at encoding by the counts of possible transitions for each lag. The actual transition is the encoding positional distance between two items at encoding (eg. recalling item 3 then item 1, transition: -1). The possible transitions is calculated from all orders in which the recalled events could have been recalled. By dividing the actual transition count with the possible transition count, a proportion is calculated for each lag. Lagged CRP was calculated for all possible transitions (-11 to 11). The list level calculation is averaged per session at each lag for a subject. This metric quantifies the degree to which items presented together are recalled together and describes the contiguity effect in each participant.

## Figures Citations

## Acknowledgments

We are grateful to the volunteers for participating in our study. We thank Srinivas Kota, PhD and Zhongzheng (Brooks) Fu, PhD for valuable feedback and all members of the Lega Lab for their contributions and patience. This project received support from National Institutes of Health grants R01-NS107357 (B.L.), and R01-NS125250 (B.L.).

---

**Figure 2—figure supplement 1.**
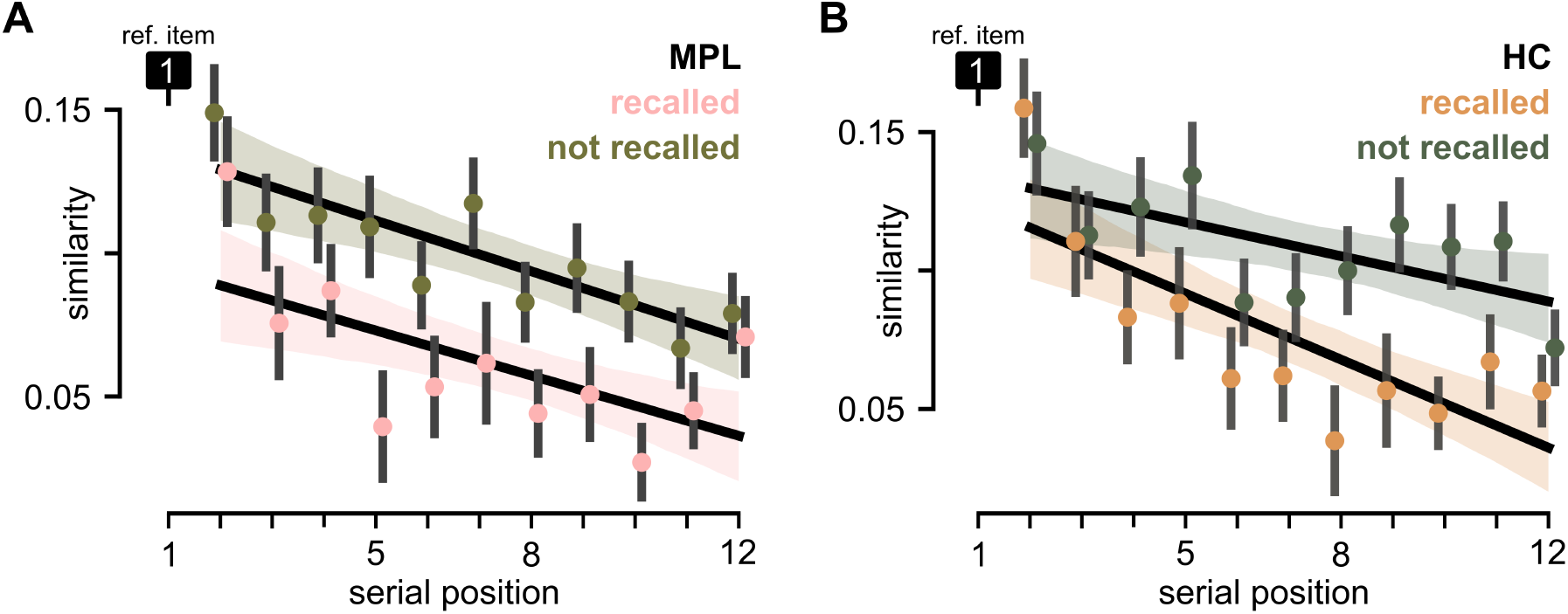
**A.** Similarity of serial position 1 to each subsequent item in the same list contrasting when both items are recalled (pink) to when both items are not recalled (green) for electrodes in the medial parietal lobe (LME, n=83; β_SP_=-0.001, CI:[-0.008, -0.003], *p*_SP_=5.6 × 10^−4^; interaction term: SP x recalled items=−9.2 × 10^−4^, CI:[-0.003, 0.004], *p*_interaction_=0.23). **B**. Same as (A) except for electrodes in the hippocampus (LME, n=76; β_SP_=0.004, CI:[-0.007, -0.001], *p*_SP_=0.004; interaction term: SP x recalled items=-0.004, [-0.008, 0.000], *p*_interaction_=0.054). All displayed trend lines are for visualization purposes only, the described LME was used to quantify the interaction effect. All error bars indicate ±1SEM.

**Figure 3—figure supplement 1.**
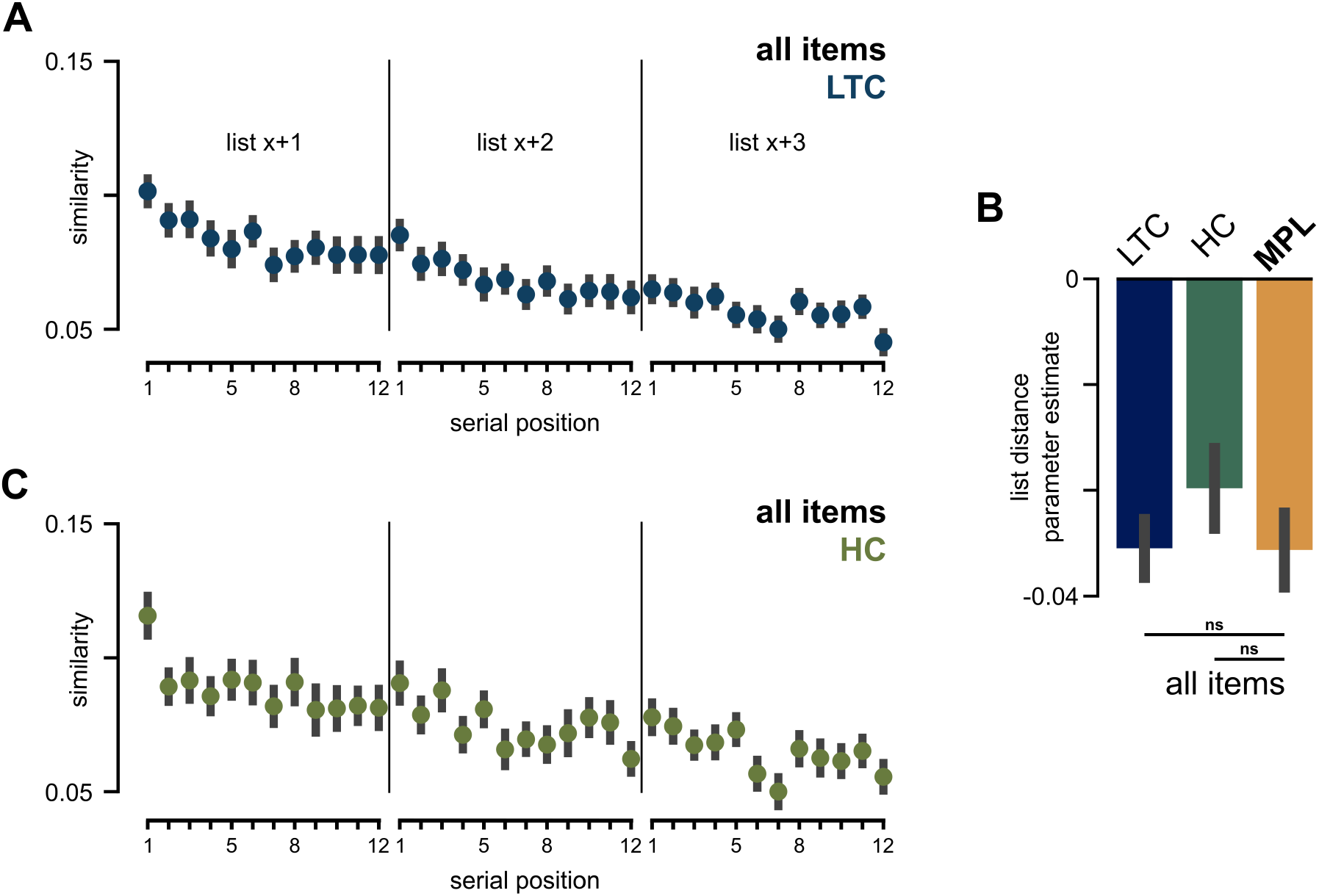
**A.** Averaged similarity of all item-pairs comparing *list x, SP1* to each item in *list x+1, x+2, x+3* for electrodes in the lateral temporal cortex. **B**. The mean parameter estimate of the predictor for list distance contrasted by region for all item pairs (t-test: MPL-LTC, t(83,81)=0.29, p=0.77; MPL-HC, t(83,76)=0.91, p=0.36 **C**. Same as (A) for electrodes in the hippocampus. All error bars indicate ±1SEM.

**Figure 3—figure supplement 2.**
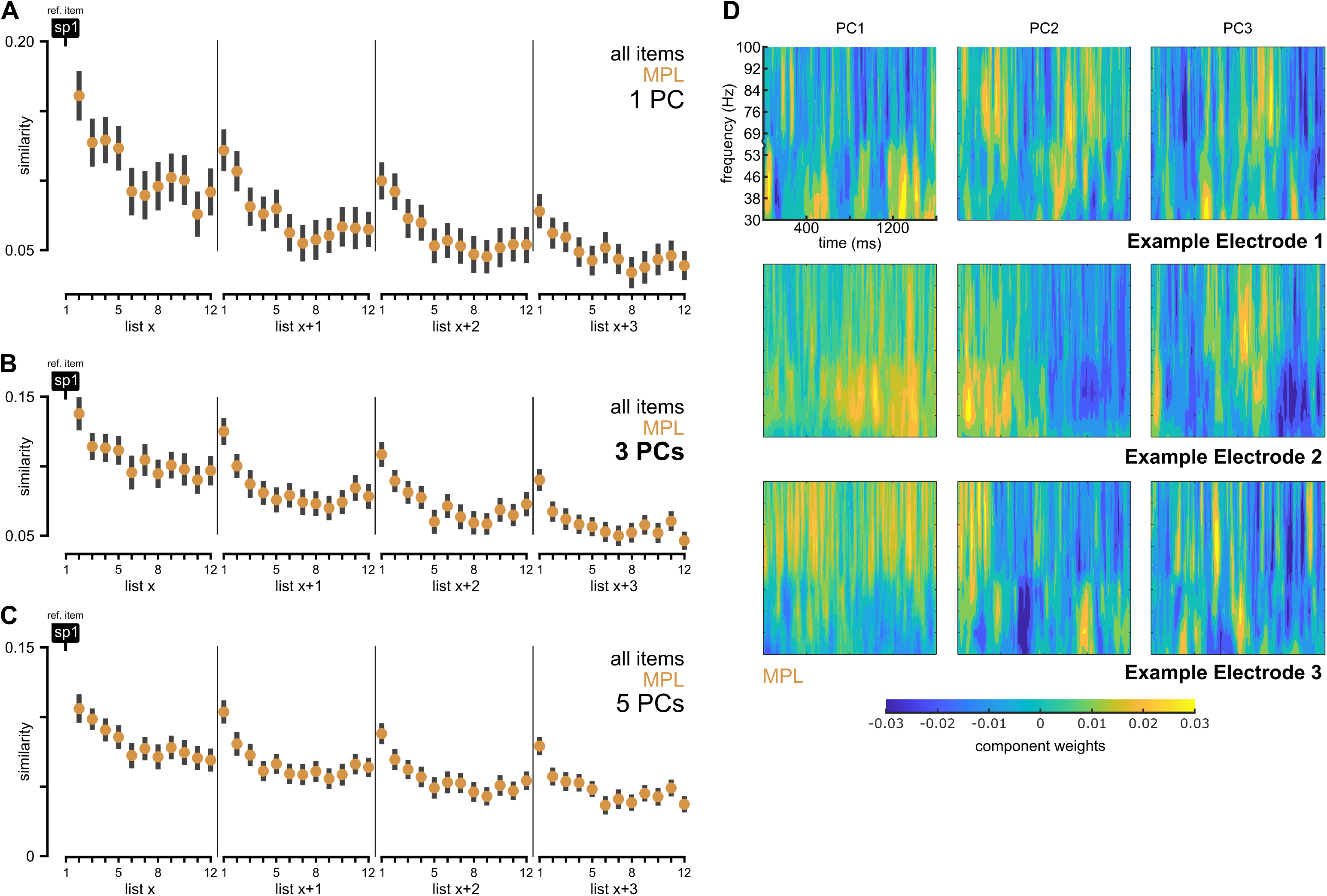
**A-C.** Averaged similarity of all item-pairs comparing *list x, SP1* to each item in *list x+1, x+2, x+3* for electrodes in the MPL. The figures demonstrate the change in average similarity when 1 principal component (PC; A), 3 PCs (B), and 5 PCs (C) are included in the similarity comparison between item time-frequency representations. **D**. Example component weighting matrices for three unique patients from the MPL. Each column represents the weighting matrix for the three eigenvalues with the highest explained variance. The matrices demonstrate unique time-frequency components underlie similarity between events. All error bars indicate ±1SEM.

**Figure 5—figure supplement 1.**
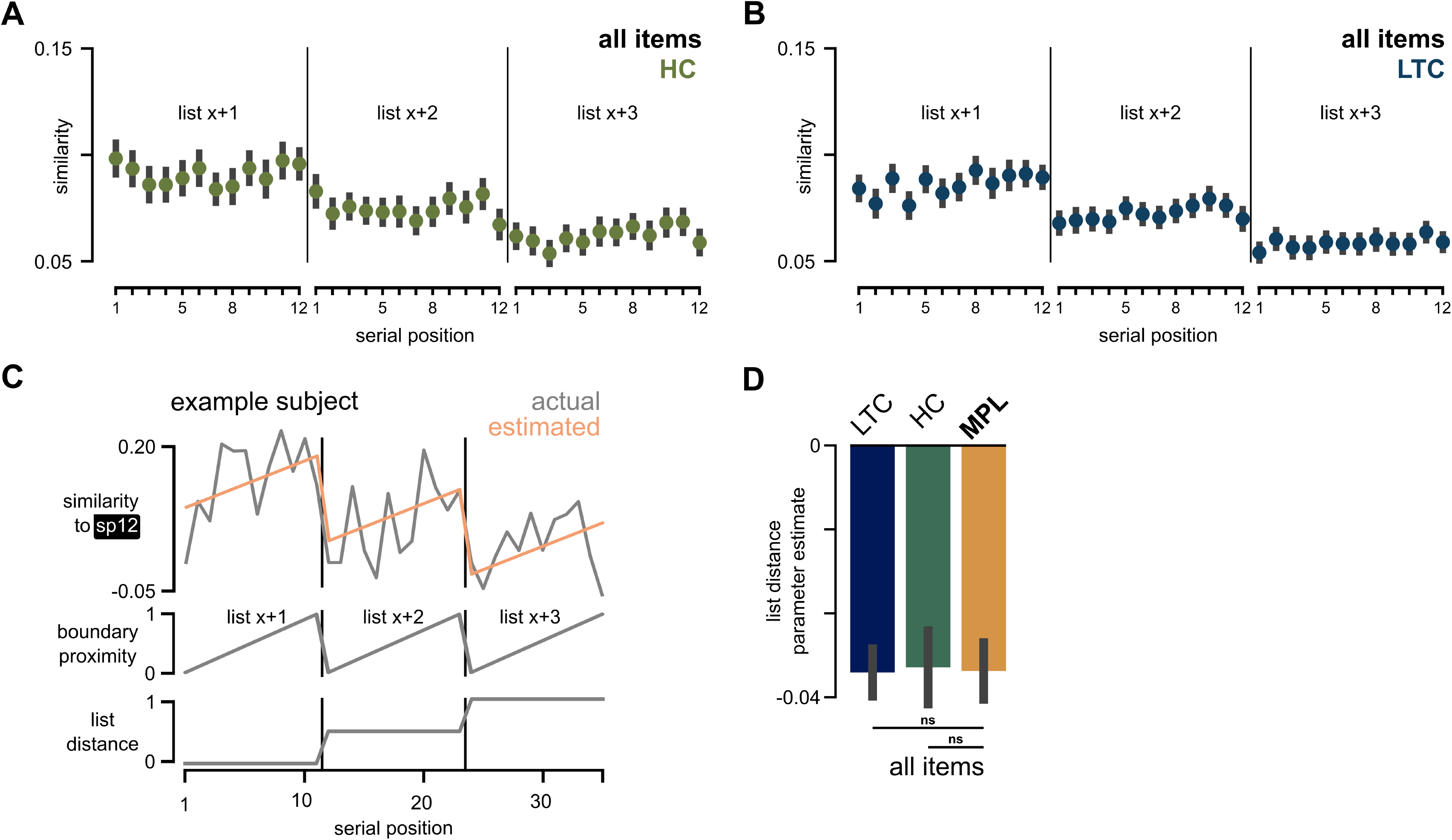
**A.** Averaged similarity of all item-pairs comparing *list x, SP12* to each item in *list x+1, x+2, x+3* for electrodes in the hippocampus. **B**. Same as (A) for electrodes in the lateral temporal cortex. **C**. Design of the GLM used to fit adjacent list similarity in the MPL for serial position 12. The response variable (similarity to *list x, SP12*; gray, top) shown for an example subject with the model prediction (orange, top) overlayed. The predictor variables, boundary proximity (middle) and list distance (bottom), are also shown. **D**. The mean parameter estimate of the predictor for list distance contrasted by region for all item pairs (t-test: MPL-LTC, t(83,81)=0.15, p=0.88; MPL-HC, t(83,76)=0.09, p=0.93. All error bars indicate ±1SEM. @par

